# Dietary non-starch polysaccharides impair immunity to enteric nematode infection

**DOI:** 10.1101/2022.07.15.500167

**Authors:** Angela H. Valente, Karen M.R. Jensen, Laura J. Myhill, Ling Zhu, Caroline M.J. Mentzel, Lukasz Krych, Henrik T. Simonsen, Josue L. Castro-Mejía, Alex Gobbi, Knud Erik Bach Knudsen, Dennis S. Nielsen, Stig M. Thamsborg, Andrew R. Williams

**Affiliations:** Department of Veterinary and Animal Sciences, University of Copenhagen, Frederiksberg, Denmark; Departmet of Food Science, University of Copenhagen, Frederiksberg, Denmark; Department of Biotechnology and Biomedicine, Technical University of Denmark, Kongens Lyngby, Denmark; Department of Plant and Environmental Sciences, University of Copenhagen, Frederiksberg, Denmark; Department of Animal Science, Aarhus University, Tjele, Denmark

**Author notes:** corresponding author –. these authors contributed equally (Angela H. Valente, Karen M.R. Jensen).

## Abstract

The influence of diet on immune function and resistance to enteric infection and disease is becoming ever more established. Highly processed, refined diets can lead to inflammation and gut microbiome dysbiosis, whilst health-promoting dietary components such as phytonutrients and fermentable fibres are thought to promote a healthy microbiome and balanced mucosal immunity. Chicory (*Cichorium intybus*) is a leafy green vegetable rich in fibres and bioactive compounds that may promote gut health. Unexpectedly, we here show that incorporation of chicory into semisynthetic AIN93G diets renders mice susceptible to infection with enteric helminths. Mice fed a high level of chicory leaves (10% dry matter) had a more diverse gut microbiota, but a diminished type-2 immune response to infection with the intestinal roundworm *Heligmosomoides polygyrus*. Furthermore, the chicory-supplemented diet significantly increased burdens of the caecum-dwelling whipworm *Trichuris muris*, concomitant with a highly skewed type-1 immune environment in caecal tissue. The chicory-supplemented diet was rich in non-starch polysaccharides, particularly uronic acids (the monomeric constituents of pectin). In accordance, mice fed pectin-supplemented AIN93G diets had higher *T. muris* burdens and reduced IgE production and expression of genes involved in type-2 immunity. Importantly, treatment of pectin-fed mice with exogenous IL-25 restored type-2 responses and was sufficient to allow *T. muris* expulsion. Collectively, our data suggest that increasing levels of fermentable, non-starch polysaccharides in refined diets compromises immunity to helminth infection in mice. This diet-infection interaction may inform new strategies for manipulating the gut environment to promote resistance to enteric parasites.

## Introduction

Diet composition may play a key role in regulation of enteric inflammation and resistance to infection [1]. The composition of human diets in affluent societies is often unbalanced, with insufficient vegetables and fruit but a surplus of easily digested carbohydrates such as starch from processed food; these conditions can drastically influence the composition of the gut microbiota (GM) [2]. The importance of a diverse and resilient GM for protection against lifestyle and infectious diseases is well established, and thus dietary components with prebiotic properties may aid in a healthy gut environment and improve immunity and disease resistance [3].

Regular consumption of green vegetables has been associated with improved immune function and less enteric pathogen infection, due to high contents of vitamins, fibres, and bioactive phytochemicals such as polyphenols [4]. Chicory (*Cichorium intybus*) is a leafy vegetable widespread across Europe and Asia, where it is grown on an industrial scale, due to its high content of the fructans (a mix of fructooligosaccharides and inulin) in the roots (up to 40% dry matter) [5]. Within the plant, the content of nutrients varies considerably; the content of inulin is negligible in the leaves, but high in the roots [6]. The health-promoting effects of inulin and its ability to alter GM composition are well known, and consequently, most studies on chicory have focused on the root part of the plant [7, 8]. However, the leaves are also widely consumed as a health-promoting food in humans reportedly having hepatoprotective and antidiabetic activity, and are traditionally used to treat diarrhoea and vomiting [9]. Recently, anti-inflammatory effects of chicory leaves in several rodent models of autoimmune inflammation have been demonstrated [10, 11].

Chicory leaves are both a rich source of fibre, which may act as prebiotic substrate for the GM, and bioactive secondary metabolites such as sesquiterpene lactones (SL) [12]. In livestock such as pigs, chicory forage consumption has been associated with higher lactobacilli:coliform ratios, most likely resulting from increased intake of soluble fibres [13, 14]. Moreover, the high concentrations of SL and other bioactive phytochemicals may reduce intestinal infections. Parasitic gastrointestinal worms (helminths) infect more than a billion people worldwide, and are also ubiquitous in livestock, and it is well known that chicory has anti-parasitic properties [15, 16]. Notably, grazing animals fed chicory have less worms [17, 18], an effect that we recently showed derives from the direct anti-parasitic properties of SL found within the plant [19]. Thus, nutraceuticals based on chicory leaves may hold great promise for reducing inflammation, promoting a healthy GM, and limiting enteric pathogens.

It is becoming increasingly apparent that there is considerable crosstalk between host dietary components and the intestinal immune system, which may have important implications for host responses to gut pathogens. Immunity to intestinal helminths is critically dependent on type-2 immune mechanisms such as the release of cytokines (IL-4, IL-13) from Th2 cells and mucus production in the gut epithelium [20]. Differences in the abundance of macronutrients (e.g. proteins), fibres, phytochemicals and vitamins may markedly affect the host mucosal response to infection [21-23]. Given the potential anti-parasitic and anti-inflammatory properties of the phytochemicals in chicory [24, 25], we hypothesized that consumption of this plant may significantly alter the host response to infection, thus providing a tractable system for dissecting the effects of phytonutrient intake on host-parasite interactions. To this end, we developed a rodent model whereby chicory leaves are incorporated into semisynthetic AIN93G rodent diets and fed to helminth-infected mice. We aimed to determine if dietary chicory could reduce helminth infection, and what impact chicory had on anti-helminth immune responses and infection-induced changes in the GM. We report here that, in contrast to chicory’s well known anti-parasitic properties in livestock, in this model chicory inclusion enhanced enteric helminth infection. We found that this unexpected result stems from non-starch polysaccharides (NSP) derived from chicory, which are lower in semisynthetic mouse diets. This increased level of NSP promoted a type-1 immune environment and susceptibility to infection, which could be reversed by exogenous administration of a type-2 cytokine. Our findings shed light on the dynamic interaction between dietary components and host immunity in the context of a pathogenic infection.

## Results

### Chicory supplementation effects immune responses to Heligmosomoides polygyrus infection, but not parasite burdens

To examine in detail how chicory influenced the course of a helminth infection in mice, we fed mice either an AIN93G control diet, or the AIN93G diet supplemented with either 1% or 10% dried chicory leaves, during infection with the small intestinal roundworm *Heligmosomoides polygyrus*. At 14 days post-infection neither faecal egg counts (FEC) or worm burdens were different between dietary treatment groups (*p*>0.05; **Figure 1A**). Infection resulted in significant eosinophilia and goblet and Paneth cell hyperplasia in the jejunum, but these parameters were unaffected by diet (**Figure 1B**). Thus, in this model dietary chicory did not exert direct anti-parasitic activity, nor influence the pathological response to infection in the small intestinal mucosa. To investigate whether chicory may influence the development of adaptive immune responses to infection, we quantified T-cell profiles in the mesenteric lymph nodes (MLN). *H. polygyrus* infection in mice fed the AIN93G diet increased the proportion of Th2 (CD4^+^ GATA3^+^) T-cells in the MLN (*p*<0.001) with no changes in the proportion of Th1 (CD4^+^ T-bet^+^) or T-regulatory (CD4^+^ Foxp3^+^) T-cells (Figure 2A). Interestingly, we noted that in infected mice fed the 10% chicory diet, there was a substantial increase (*p* < 0.01 for interaction between diet and infection) in Th1 cells following infection that did not occur in the other groups, together with a small, non-significant decrease in Th2 cells (**Figure 2A**). Thus, the Th2: Th1 ratio in infected mice fed the 10% chicory diet was significantly skewed towards a Th1 profile compared to control fed-infected mice, which had a highly polarized Th2 profile characteristic of helminth infection (**Figure 2B**). There was no effect of diet on T-regulatory cell proportions in either uninfected or *H. polygyrus*-infected mice (**Figure 2A**). To confirm further the Th1 polarization, we performed qPCR on tissue from the proximal small intestine to investigate the expression of *Dclk, Duox2, Ifng, Il10, and Gpx2*. Gene expression confirmed the chicory-mediated polarization towards Th1 cells revealed by T-cell phenotyping. Expression of the tuft cell marker *Dckl1*, which was highly induced by *H. polygyrus* infection, was significantly suppressed by chicory in a dose-dependent manner **(Figure 2C**). Similarly, infection-induced expression of *Duox2* and *Gpx2* was also attenuated by dietary chicory, albeit not significant (**Figure 2C**). In contrast, expression of the Th1/Treg related genes *Ifng* and *Il10* was either unaffected or tended to be increased by chicory supplementation (**Figure 2C**). Collectively, these data show that chicory did not lower *H. polygyrus* infection, and, in fact, appeared to promote a Th1 immune response which is normally associated with helminth persistence.

**Figure 1.**
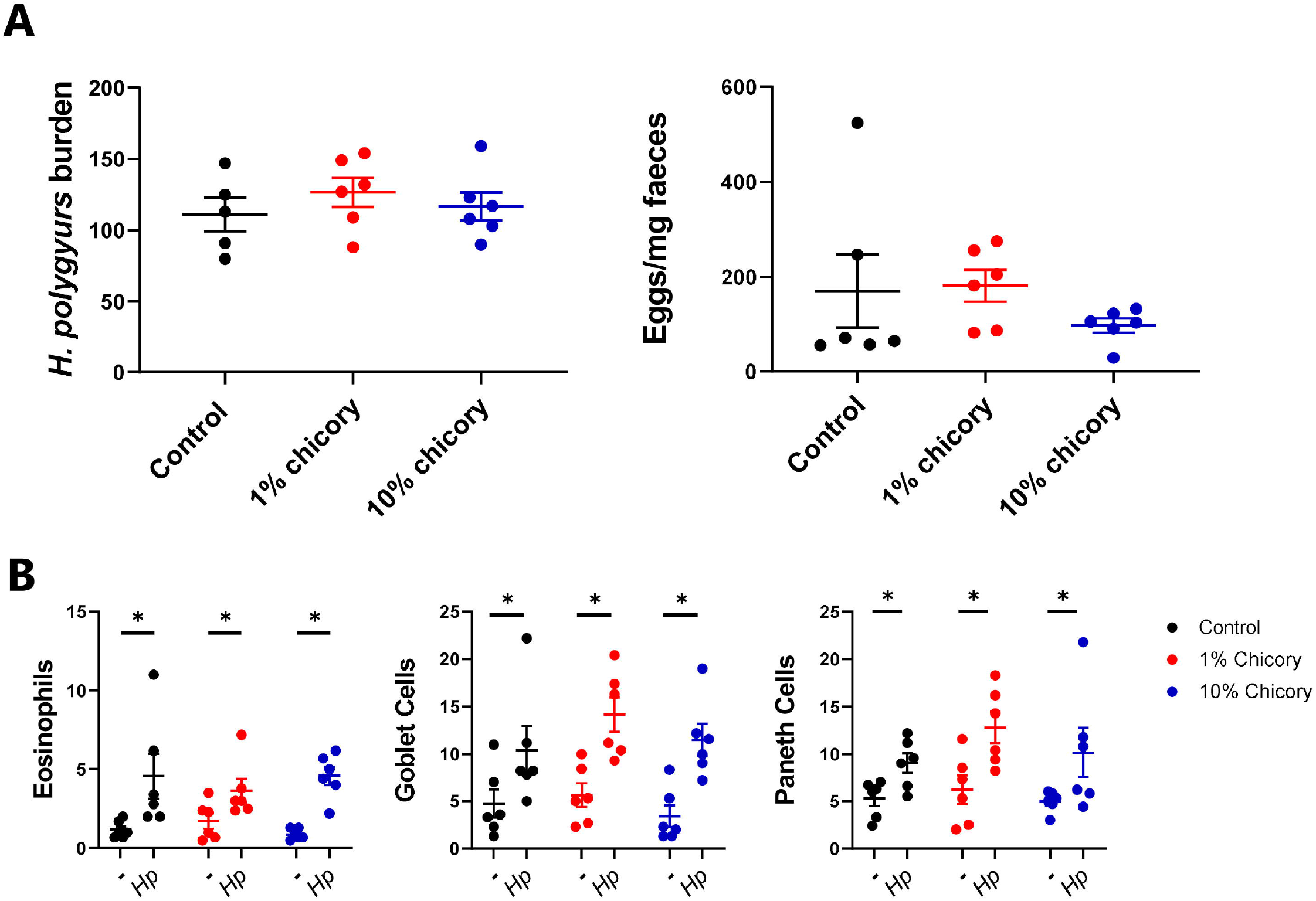
Dietary chicory does not affect *Heligmosomoides polygyrus* burdens or histopathological responses. **A)** *H. polygyrus* worm burdens and faecal egg counts 14 days post-infection, in mice fed either control AIN93G diets or the control diet supplemented with chicory (1% or 10% dry matter). **B)** Eosinophil, goblet cell and Paneth cell responses in jejunal tissue (cells/mm tissue). n= 6 per group, \**p*<0.05 by ANOVA.

**Figure 2.**
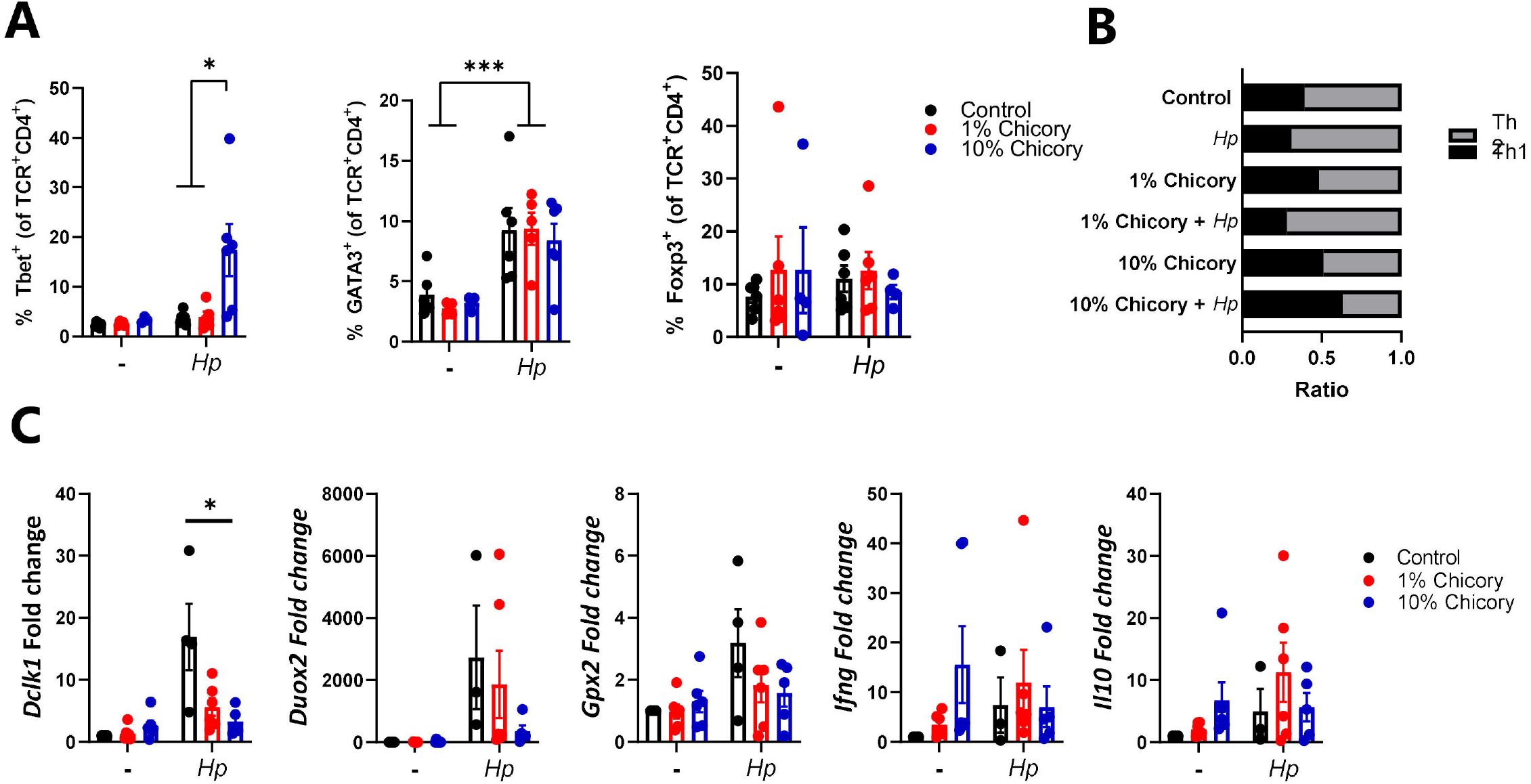
Chicory modulates the immune response to *Heligmosomoides polygyrus* infection. **A)** Percentage of CD4 Tbet CD4 GATA3 and CD4 Foxp3 T-cells in the mesenteric lymph nodes 14 days post-infection in mice fed either control AIN93G diets or the control diet supplemented with chicory (1% or 10% dry matter). **B)** ratio of CD4 GATA3 /CD4 Tbet T-cells in lymph nodes. **C)** Expression of selected genes in jejunum tissue. *p<0.05; ***p<0.001 by ANOVA, n=5-6 per group.

### Dietary chicory increases diversity and modulates Heligmosomoides polygyrus-induced changes in the gut microbiota

Chicory contains a number of putative prebiotic constituents that may modify the host GM [26]. To determine if the chicory-mediated changes in immune function were accompanied by changes in the GM, caecal digesta samples were analysed by 16S rRNA gene amplicon-based sequencing. Interestingly, the mice fed the 10% chicory diet had a significantly more diverse GM than the other groups **(Figure 3A)**. Furthermore, samples from the three dietary groups clustered distinctly based on unweighted UniFrac distance metrics (**Figure 4B**). Samples from mice fed the 10% chicory diet were most divergent from mice fed the AIN93G control diet, with mice fed the 1% chicory diet clustering closer with controls (intermediate) (**Figure 3B;** *p*<0.05 by PERMANOVA). Uninfected and *H. polygyrus*-infected mice also clustered into two distinct groups (**Figure 3C**; *p*<0.05 by PERMANOVA). Notably, when plotting Unweighted UniFrac distance metrics based on the 3×2 factorial design showed 6 distinct clusters, indicating profound interactions between the treatments and indicating that chicory modulated the *H. polygyrus*-induced changes in the GM (**Figure 3D**).

**Figure 3.**
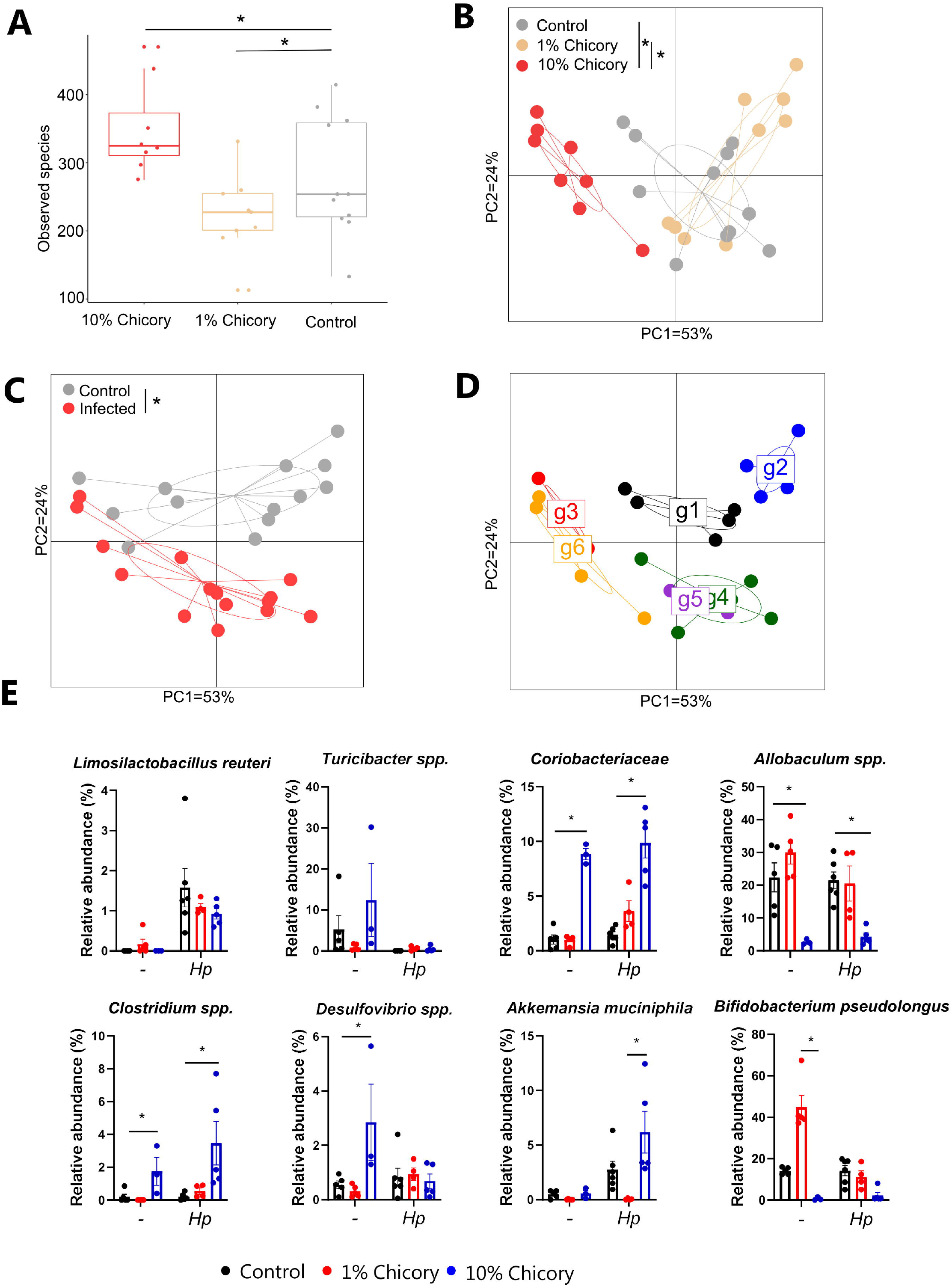
Effects of chicory and *Heligmosomoides polygyrus* infection on composition of the cecal microbiota. **(A)** α-diversity box plots showing higher diversity in mice fed 10% chicory. **(B-D)** β-diversity by unweighted UniFrac showing a divergence in mice with different treatments. **(E)** Abundance of taxa identified by ANCOM as being significantly impacted by diet and/or infection. Differentially abundant taxa were further analysed using Kruskal-Wallis testing within each infection group. n=5-6 mice per group. * p < 0.05 by Kruskal Wallis-test.

**Figure 4.**
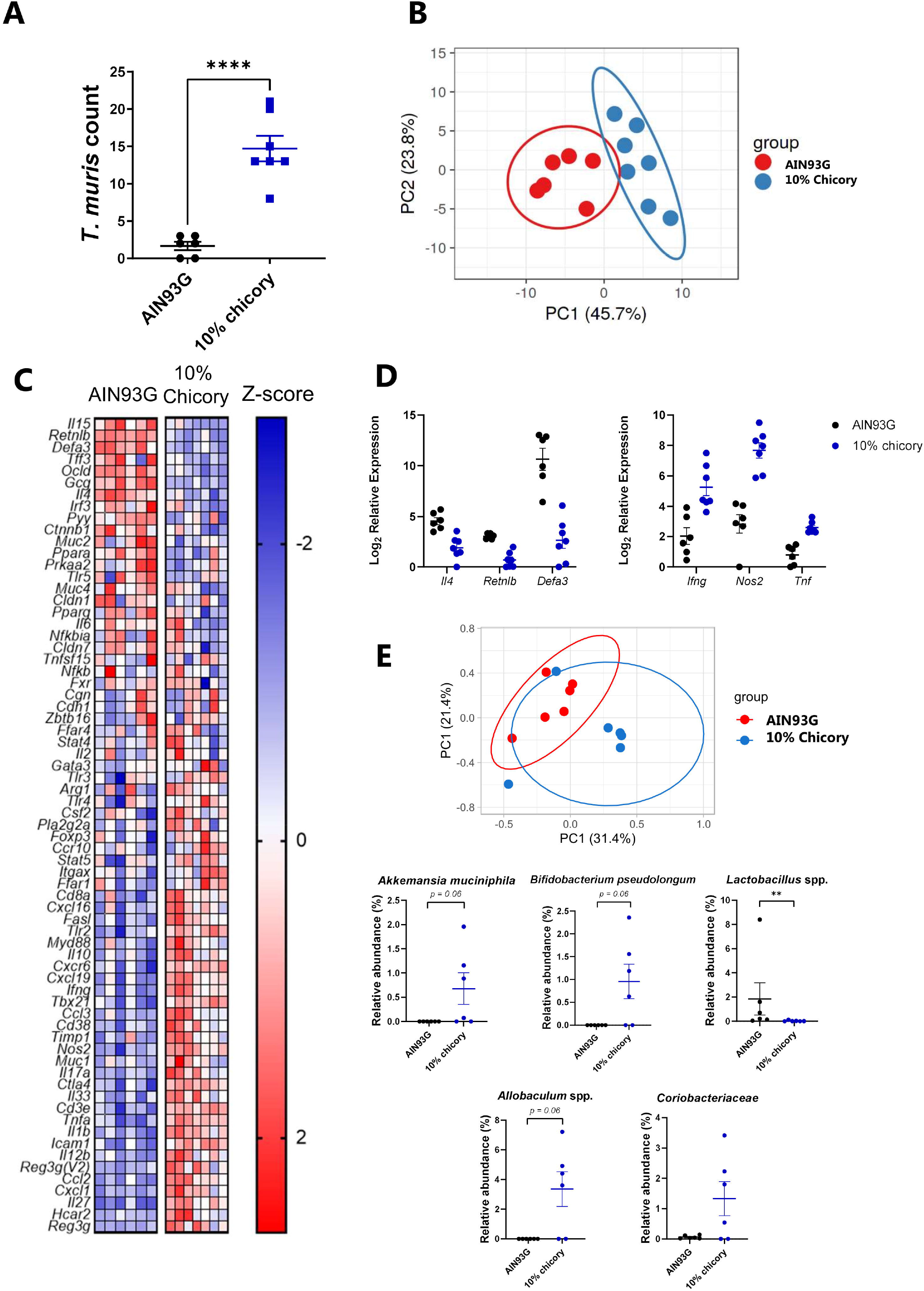
Dietary chicory increases *Trichuris muris* burdens and alters immune responses and gut microbiota composition. **A)** Worm burdens at day 35 after the start of trickle infection in mice fed either a control AIN93G diet or the control diet supplemented with 10% chicory. Mice were infected with 20 eggs at day 0, 7 and 14. n=6-7 mice per group, **** *p* < 0.0001 by T-test. **B)** Principal Component Analysis plots showing clustering of groups based on Fluidigm gene expression analysis. **C)** Heat map of gene expression in caecal tissue. **D**) Mean relative expression of selected genes in caecal tissue. **E)** Principal Coordinates Analysis plots showing clustering of groups based on 16S rRNA gene amplicon based sequencing (Bray-Curtis Dissimilarity Metrics) of caecal microbiota and relative abundance of zOTUs identified as being significantly impacted by diet by ANCOM analysis (followed by Mann-Whitney testing). n=6 mice per group. ** *p* < 0.01 by Mann-Whitney test.

ANCOM analysis identified 8 bacterial taxa that were significantly differently distributed between the groups (**Figure 3E**). Independently of diet, *H. polygyrus* infection decreased the abundance of *Turicibacter spp*. and increased the abundance of a zOTU (zero-radius Operational Taxonomic Unit) closely related to *Limosilactobacillus reuteri* (**Figure 3E**). The effects of chicory were markedly dependent on the inclusion level in the diet. Ten % chicory had profound effects on the relative abundance of *Coriobacteriaea* and *Clostridia* spp. (both increased) and *Allobaculum spp*. (decreased), but these effects were not evident with 1% chicory. Significant interactions were observed between diet and infection for the abundances of Desulfovibrio spp. and zOTUs closely related to *Akkermansia muciniphila* and *Bifidobacterium pseudolongum*. Ten % chicory tended to increase the abundance of *Desulfovibrio spp*. in uninfected mice, but decreased it in infected mice. The abundance of *A. muciniphila* was increased by infection in control-fed mice and those fed 10% chicory, but not those fed 1% chicory. Finally, the abundance of *B. pseduolongum* was markedly increased by 1% chicory, particularly in uninfected mice, but was strongly suppressed by 10% chicory. Collectively, these data indicate that supplementing an AIN93G diet with 10% chicory substantially increases the α-diversity and abundance of specific/several zOTUs, and significantly modulates *H. polygyrus -induced change*s, in the caecal GM.

### Dietary chicory increases Trichuris muris burdens in the caecum and creates a polarized type-1 environment compared to AIN93G diets

As we observed an effect of chicory on *H. polygyrus*-induced immune responses and GM composition, but not on worm burdens, we next asked whether chicory may have a stronger modulatory effect on a caecum-dwelling parasite, at the main site of microbial fermentation in the gut. As the 10% chicory diet induced the strongest effect on immune function and the GM, we tested the effect of this diet in mice infected with the whipworm *T. muris*. Mice were trickle-infected with three doses of 20 *T. muris* eggs, which stimulates a chronic infection in C57BL/6 mice [27, 28]. Strikingly, five weeks after the commencement of trickle infection, mice fed the chicory-supplemented diet had significantly higher worm burdens than mice fed the AIN93G diet (**Figure 4A)**.

To explore the immunological basis underlying the effect of chicory on *T. muris* burdens, we performed high-throughput Fluidigm-based qPCR to measure expression of a panel of immune and mucosal barrier-related genes in caecum tissue of infected mice. Mice fed either AIN93G or the 10% chicory diet clustered into two distinct groups based on their transcriptional response (**Figure 4B**). We observed a broad down-regulation of genes involved in Th2 immune function and mucosal barrier defences, and a strong upregulation of genes involved in Th1 and Th17 immune function (**Figure 4C**). Genes that were significantly upregulated in the mice fed chicory included *Ifng, Nos2 a*nd *Tnf*, whilst downregulated genes involved *Il4, Retnlb* and *Defa3* (**Figure 4D**). The caecal microbiota composition differed between the dietary groups (*p*=0.055 based on Bray-Curtis Dissimilarity metrics; **Figure 4E**), but not on unweighted UniFrac distance metrics (*p*=0.25; data not shown). We noted that, consistent with the experiments in *H. polygyrus*-infected mice, chicory caused an expansion of *A. muciniphila*, and the *Coriobacteriaceae* family (**Figure 4E**). In contrast to *H. polygyrus*-infected mice, in this experiment the abundance of *Allobaculum spp*. and zOTUs corresponding to *B. pseudolongum* were higher in chicory-fed mice. Notably, mice fed the chicory-supplemented diet had significantly lower levels of *Lactobacillus* spp (**Figure 4E**). Taken together, these data indicate that inclusion of 10% chicory into a purified diet altered the gut environment and promoted *T. muris* infection, by inhibition of protective type-2 immune mechanisms.

### Dietary pectin promotes Trichuris muris infection and impairs type-2 immunity

To investigate the underlying cause of the impaired type-2 response and increased *T. muris* burdens in chicory-fed mice, we first performed a detailed chemical composition of the different diets. Chemical analysis showed that the diets with chicory inclusion (particularly at 10% inclusion) were enriched in dietary fibre, especially NSP, compared to the control diet (Table 1). The 10% chicory diet contained twice the amount of non-cellulosic, NSP as the purified AIN93G diet, including the presence of arabinose, galactose and uronic acids that were absent in the control diet. This prompted us to examine the role of NSP in immunity to *T. muris*. Uronic acids are the monomeric backbone of pectin, a major component of the primary cell walls of plants, and a prebiotic source of fermentable fibre for the GM [13, 29]. To determine if an increased pectin content could be responsible for the observed effects on anti-helminth immunity, we fed mice either an AIN93G diet or the same diet supplemented with 5% pectin during a trickle *T. muris* infection i.e. 3 doses of 20 eggs weekly. Notably, pectin-fed mice had significantly higher worm burdens, along with substantially higher serum levels of *T. muris*-specific IgG2a (a marker of a Th1 response; **Figure 5A**). Furthermore, pectin-fed mice had increased levels of *Ifng* and *Nos2* expression, and decreased levels of *Retnlb and Il13* expression, in caecal tissue (**Figure 5B**). Thus, dietary pectin supplementation closely resembled the observed effects of chicory in impairing immunity to *T. muris*. To further demonstrate a role for pectin in inhibiting the type-2 response, we fed mice either AIN93G or pectin-supplemented diets and infected them with a single dose of 300 *T. muris* eggs, which typically stimulates a highly skewed type-2 response and worm expulsion around 21 days post-infection (p.i.) [30]. Indeed, mice fed the AIN93G diet had largely cleared their infection at day 21 p.i., whereas pectin-fed mice all still harboured worms, indicating expulsion was incomplete (Figure 5C). Consistent with this, levels of *T. muris*-specific IgE were significantly lower in pectin-fed mice, whereas IgG1 and IgG2a levels were unaffected (Figure 5D). Thus, inclusion of a non-cellulosic NSP source is sufficient to impair type-2 immunity and *T. muris* expulsion.

**Table 1.** Composition of Experimental Diets

|  | AIN93G | 1% chicory | 10% chicory | 5% Pectin |
| --- | --- | --- | --- | --- |
| ***Ingredients (g/100g)*** |  |  |  |  |
| Casein | 20 | 19.9 | 18.96 | 20 |
| L-cysteine | 0.3 | 0.3 | 0.3 | 0.3 |
| Corn Starch | 39.75 | 38.83 | 30.75 | 34.75 |
| Maltodextrin | 13.2 | 13.2 | 13.2 | 13.2 |
| Sucrose | 10 | 10 | 10 | 10 |
| Cellulose | 5 | 5 | 5 | 5 |
| Chicory Leaves<sup>1</sup> | 0 | 1 | 10 | 0 |
| Pectin<sup>2</sup> | 0 | 0 | 0 | 5 |
| Vitamin premix | 1 | 1 | 1 | 1 |
| Mineral premix | 3.5 | 3.5 | 3.5 | 3.5 |
| Choline bitartrate | 0.25 | 0.25 | 0.25 | 0.25 |
| TBHQ | 0.0014 | 0.0014 | 0.0014 | 0.0014 |
| Soybean Oil | 7 | 7 | 7 | 7 |
| ***Calculated Composition (%)*** |  |  |  |  |
| Crude Protein | 17.6 | 17.6 | 17.6 | 17.6 |
| Crude Fat | 7.1 | 7.1 | 7.1 | 7.1 |
| **Energy (MJ ME/kg)** | 16.2 | 16.2 | 15.8 | 15.6 |
| ***Analyzed Composition (%)*** |  |  |  |  |
| Glucose and Fructose | 0.05 | 0.1 | 0.55 | ND |
| Sucrose | 13.28 | 13.17 | 14 | ND |
| Fructans | 0 | 0 | 0 | ND |
| Starch | 54.85 | 51.37 | 47.64 | ND |
| **Non-cellulosic polysaccharides (%)** |  |  |  |  |
| Rhamnose | 0 | 0 | 0 | ND |
| Fucose | 0 | 0 | 0 | ND |
| Arabinose | 0 | 0 | 0.2 | ND |
| Xylose | 1.1 | 1.0 | 1.2 | ND |
| Mannose | 0.1 | 0.1 | 0.2 | ND |
| Galactose | 0 | 0 | 0.3 | ND |
| Glucose | 0.4 | 0.5 | 0.4 | ND |
| Uronic Acids | 0 | 0.2 | 1.2 | ND |
| Total Non-cellulosic polysaccharides | 1.6 | 1.8 | 3.5 | ND |
| Cellulose | 3.7 | 3.6 | 4.7 | ND |
| Lignin | 0.3 | 0.3 | 0.3 | ND |
| **Total dietary fibre (%)** | 5.6 | 5.7 | 8.5 | ND |

**Figure 5.**
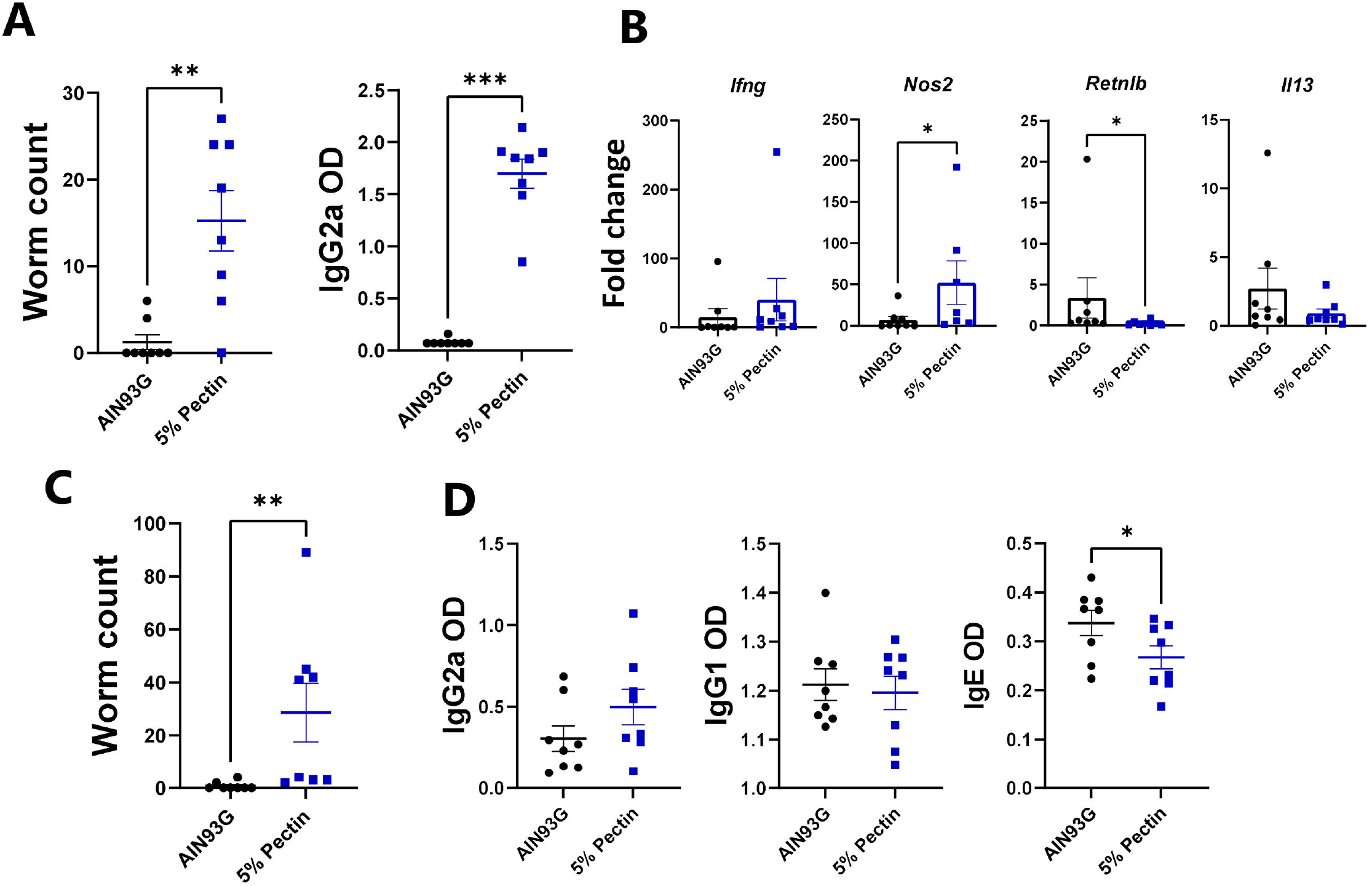
Dietary pectin increases *Trichuris muris* burdens and impairs type-2 immune responses. **A)** Worm burdens at day 35 after the start of trickle infection and *T. muris*-specific serum antibody levels in mice fed either a control AIN93G diet or the control diet supplemented with 5% pectin. Mice were infected with 20 eggs at day 0, 7 and 14. **B)** Expression of selected genes in caecal tissue, shown as fold change, **C**) Worm burdens, and **D)** antibody responses in mice 21 days post-infection with 300 *T. muris* eggs. n= 8 mice per group. * *p* <0.05; ** *p*<0.01; \*\**p*<0.001 0.001 by t-test or Mann-Whitney test.

### IL-25 treatment restores immunity to Trichuris muris in pectin-fed mice

We postulated that the inclusion of dietary NSP impaired worm expulsion by promoting a type-1 immune response at the expense of type-2 immunity. However, the possibility remained that chicory or pectin may increase worm burdens by an alternative mechanism, e.g. by acting as an additional nutrient source for the parasites. We reasoned that if impaired immunity was responsible, then exogenous addition of a type-2 polarising factor should overcome the dietary factors to promote expulsion. IL-25 is an alarmin released by epithelial cells during allergies and helminth infection, which can drive a multi-faceted type-2 response and restore immunity in susceptible mice (e.g. SCID mice) when administered exogenously [31]. Therefore, mice were fed pectin-supplemented diets during a high-dose *T. muris* infection, with or without IL-25 treatment. Consistent with previous data, mice fed an AIN93G diet had cleared the infection at day 21 p.i., whereas expulsion was incomplete in pectin-fed mice administered vehicle control (**Figure 6A**). Importantly, IL-25 treatment completely restored expulsion (**Figure 6A**). Measurement of serum antibodies showed that rIL-25 treatment boosted IgE responses that were impaired by dietary pectin (**Figure 6B**), whilst analysis of MLN cells showed that pectin diminished Th2 (GATA3) T-helper responses, but IL-25 treatment effectively restored a Th2 dominance within the MLN (**Figure 6C**). Thus, strengthening the host type-2 response was sufficient to abrogate the effect of the dietary pectin, suggesting that the role of dietary NSP in impairing anti-helminth immunity is by skewing the type-1/type-2 response in favour of a type-1 environment which promotes worm survival.

**Figure 6.**
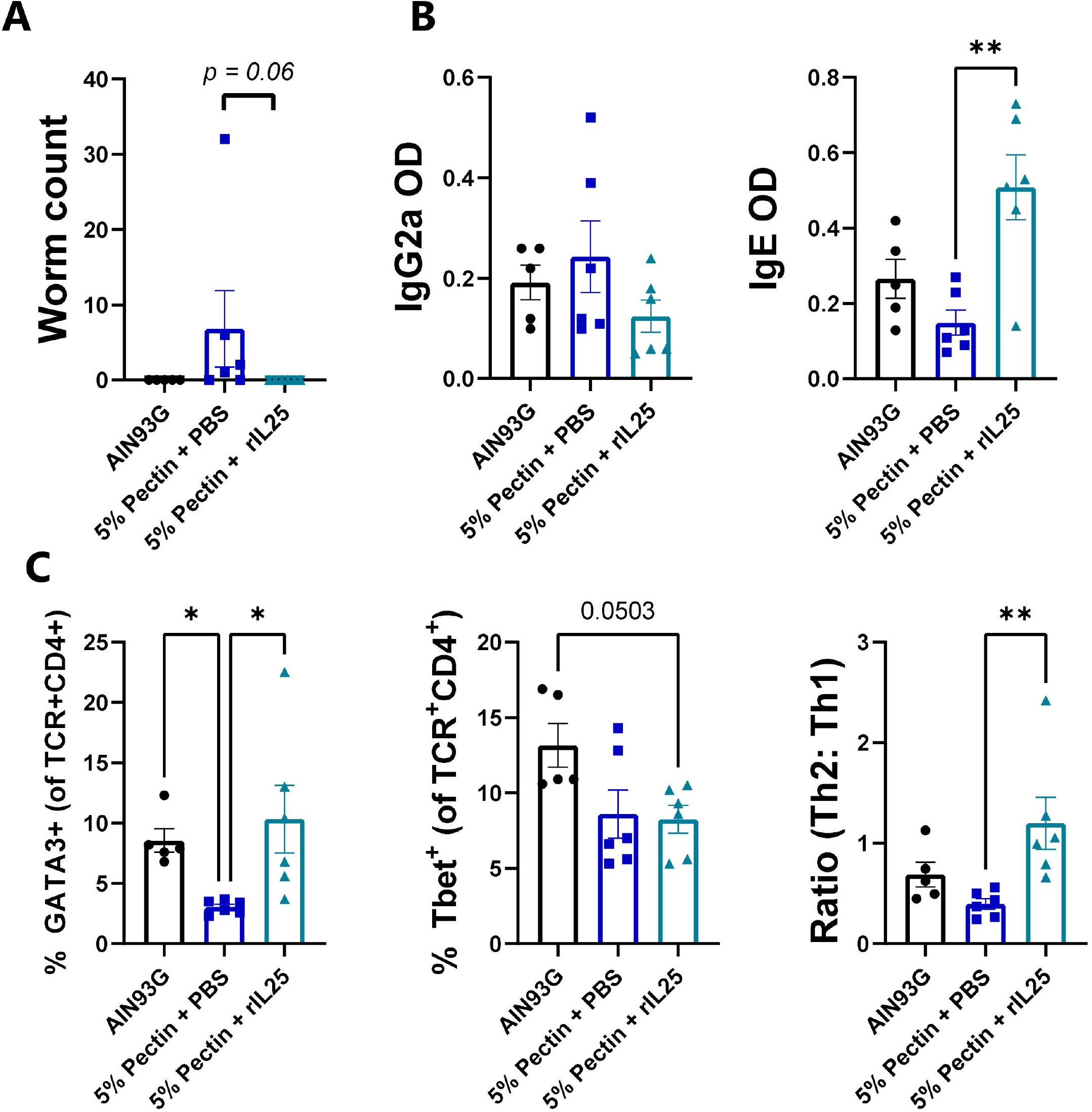
IL-25 treatment restores *Trichuris muris* expulsion in mice fed pectin. **A)** Worm burdens 21 days post-infection with 300 *T. muris* eggs in mice fed an AIN93G control diet, or the control diet supplemented with 5% pectin and administered either PBS or rIL-25 every three days from day 5 post-infection to day 18 post-infection, **B)** *T. muris*-specific serum antibody levels and **C)** T-cell proportions in mesenteric lymph nodes at day 21 post-infection. n = 6 mice per group. \**p*<0.05; \*\**p*<0.01 by one-way ANOVA or Kruskal-Wallis test.

## Discussion

Bioactive dietary components play a large role in regulating gut health and resistance to enteric pathogen infections. Our recent work with chicory has identified that SL have a clear anthelmintic activity, and it is known that SL (especially those derived from chicory) are anti-inflammatory and antibacterial compounds [32, 33]. Thus, we examined the effects on parasitic infection and gut microbiota in mice. Unexpectedly, rather than reducing parasite burdens, we found that chicory increased whipworm infection, and in both *T. muris* and *H. polygyrus* models, inclusion of chicory in the diet resulted in a significant muting of the helminth type-2 response. Thus, in our mouse model, it may be possible that an accompanying high fiber content overruled the potential anthelmintic effects we expected to find due to the SL content of the chicory in the diet.

Incorporation of chicory into AIN93G diets was done so as to keep the diets iso-caloric and balanced for crude protein content, but the diets thus differed in their fibre content, suggesting a possible mechanism for the observed effects. Whilst cellulose content was higher in the chicory-supplemented diets, we have previously shown that cellulose content in the diet does not result in increased *T. muris* infections [30], suggesting that the most likely cause was the increase in non-cellulosic polysaccharides in the chicory diet; the latter was confirmed by experiments with pure pectin substituting for chicory. Whilst the difference in non-cellulosic polysaccharide content between the AIN93G diet and the 10% chicory-supplemented diet was marginal (an increase from 1.8% to 3.6%), it was clearly sufficient to have a strong effect on *T. muris* infection. We cannot rule out that other NSP sugar residues such as galactose may also play a role in this response, or that they may be synergistic effects of different NSP within complex plant material in regulating the response to helminth infection.

The exact mechanisms whereby dietary NSP impair type-2 immunity remains to be established, but it is telling that in *H. polygyrus*-infected mice, high levels of chicory caused increased α-diversity in the GM. A species-rich GM may produce a plethora of metabolites that influence the activity of immune cells [34]. For example, species richness has been correlated with the production of metabolites such as secondary bile acids which promote the production of type-17 cytokines such as IL-22, but may potentially inhibit the expression of type-2 anti-helminth immunity [35, 36]. Consistent with this hypothesis, germ-free mice have markedly higher Th2 responses than conventional mice and are refractory to infection with *T. muris* and also somewhat more resistant to *H. polygyrus* [37, 38]. This may suggest that an increasingly rich GM may progressively shift the mucosal immune response towards more of a type-1 or type-17 environment and downplay Th2 responses. More studies are needed to unravel these mechanisms.

Interestingly, we have recently shown that semisynthetic mouse diets supplemented with pure inulin, a highly fermentable fructan polymer, also impaired type-2 immune responses to *T. muris* and prevented worm expulsion [30]. This may suggest that a range of prebiotic substrates, including both highly refined polymers such as inulin and pectin, and crude plant material such as chicory leaves, can modulate the response to helminths in a negative way. However, we have also noted, that including inulin in a large animal model of helminth infection (*Trichuris suis* in pigs) did not have a type-1 polarizing effect, and in fact tended to promote Th2 responses [39]. The reasons for these discrepancies between studies have not yet been resolved, but may be related to differences in diet composition between the semisynthetic nature of compositionally defined mouse diets and the complex nature of pig diets which moderates the effects of the inulin inclusion due to a pre-existing and higher level of fermentable fiber in the basal diet. Consistent with this, it has in some mouse models been shown that colitis can be worsened by inclusion of high levels of inulin, but only when incorporated into semisynthetic diets and not unrefined mouse chow [40]. Thus, there appears to be a complex trilateral relationship between diet, the GM and mucosal immune function during pathogen infection. Elucidating the underlying mechanisms of this interaction would be highly valuable for the development of targeted nutritional interventions to fine-tune immune response in the context of different gut infections.

In conclusion, we have shown here a novel role for plant NSP in mediating susceptibility to enteric helminth infection. An increased understanding of the factors underlying this effect of diet may allow development of new tools to manipulate the gut environment to promote resistance to enteric pathogens.

## Materials & methods

### Mice

All animal experimentation was conducted under the guidelines and with approval of the Danish Animal Experimentation Inspectorate (Licence number 2015-15-0201-00760). In all experiments female C57BL/6 mice (aged 6-8 weeks; Envigo) were used. Mice were kept in individually-ventilated cages with sawdust, nesting material and *ad libitum* water and feed. In all experiments, mice were allowed to adapt to their respective diets for two weeks prior to infection,.

### Parasite Infection, IL-25 treatment, and Necropsy

*H. polygyrus* and *T. muris* were propagated as previous described [30, 41]. For *H. polygyrus* infection, mice were infected with a single dose of 200 L3 by oral gavage and killed 14 days p.i. For *T. muris* infection, mice received a trickle infection consisting of 20 eggs by oral gavage at days 0, 7 and 14 before sacrifice at day 35 post first infection dose, or a single dose of 300 eggs followed by sacrifice at day 21 p.i.. Where indicated, mice received IL-25 treatment during high-dose (300 eggs) *T. muris* infection by i.p. injection of 5 µg rIL-25 (Biolegend #587302), or vehicle control (PBS), on days 5,8,11,14 and 17 p.i. All mice were sacrificed by cervical dislocation. Immediately after termination, approximately 0.5 cm of the proximal jejunum and/or caecum was collected and stored in RNAlater (Sigma-Aldrich). For *H. polygyrus*-infected mice, an additional 1 cm cut from the proximal end of the jejunum was collected and stored in 10% natural buffered formalin (4% formaldehyde) for histology, and cell numbers enumerated by a microscopist (blinded to the treatment groups). Fresh digesta samples were collected from the caecum and snap frozen at -80 °C for GM analyses. Furthermore, the mesenteric lymph nodes (MLN) were collected and stored in RPMI 1640 media (Sigma-Aldrich). The MLNs were supplemented with 10% fetal calf serum (Sigma-Aldrich) and stored on ice for flow cytometry (see below). Worm count was performed by manually picking as the mice were euthanized and faeces taken from the colon was stored for egg count (FEC) by the modified McMaster technique [42].

### Experimental diets

The chicory used for the experimental diets was *C. intybus* cv. “Spadona” (DSV Ltd., Denmark) sown as a pure sward (7 kg seeds/ha) in May 2019 and harvested mid-June 2019 at the experimental facilities of the University of Copenhagen (Taastrup, Denmark, 55_6704800N, 12_2907500E). Approximately 12 kg of fresh chicory leaves were harvested and dried using a constant airflow. The dry and ground leaves were incorporated into a purified AIN93G diet at the expense of starch and casein, with balanced crude protein and metabolisable energy content, and processed into pellets. The pelleting procedure did not involve heating above 30°C. Three diets were prepared (Table 1): A control diet (standard AIN93G), and two experimental diets with 1% and 10% chicory (Table 1). In addition, a diet with 5% citrus peel pectin (Sigma-Aldrich) in place of starch was also formulated (Table 1) using the same pelleting procedure. All diets were prepared by Sniff Spezialdiäten GmbH, Germany.

### Flow cytometry

MLNs were processed using a 70 μM cell strainer. The cell suspension was then washed and suspended at 5 × 10 cells/mL. Before staining, cells were washed in cold PBA with 2% FCS, and Fc receptors were blocked using FC Block (1:100) (BD biosciences # 553141). Cells were then stained in 96-well round-bottom plates. For extracellular staining, the cells were stained with TCRβ-FITC (clone H57-597; BD Biosciences) and CD4–PerCP-Cy5.5 (clone R4-5; BD Biosciences). For intercellular staining, cells were permeabilized using the FoxP3/ transcription factor staining buffer set (Thermo Fisher) and then incubated with the following antibodies: Tbet-AlexaFluor 647 clone (4B10; BD Biosciences), GATA3–PE-conjugated (clone TWAJ; ThermoFisher) or FoxP3–FITC (clone FJK-16s; ThermoFisher). FMO and isotype controls were included. After staining cells were processed using a BD Accuri C6 flow cytometer (BD Biosciences) and data was acquired and analysed using Accuri CFlowPlus software (Accuri® Cytometers Inc., MI, USA).

### 16S rRNA gene Amplicon Sequencing

DNA was extracted from caecum digesta using a commercial Bead-Beat Micro AX Gravity kit (A&A Biotechnology), used in accordance with manufacture’s guidelines. Before extraction, samples were lysed in lysis buffer supplemented with lysozyme (4000 U) and mutanolysin (50 U) and incubated at 50°C for 20 min. DNA concentration was measured on Varioskan® (Thermo Fisher Scientific). The V3 region on the 16S rRNA gene was amplified using the universal forward primer 338 F (5’-ACTCCTACGGGAGGCAGCAG-3’) and reverse primer 518 R (3’-ATTACCGCGGCTGCTGG-5’) (0.5 μL of each/sample, at 10 μM concentration) that included Nextera™ (illumina CA, USA) compatible overhangs. In addition to this, 5 μL/sample 5x PCRBIO HiFi buffer (PCR Biosystems©, UK), 0.25 μL/sample PCRBIO HiFI Polymerase (PCR Biosystems ©, UK), and 1 μL/sample BSA buffer (Sigma) and formamide was added. 17.75 μL DNA was added after being diluted to 1/500 using sterilized Milli-Q® water, resulting in a total volume of 25 μL. Standard PCR cycling was used: One initial 95 °C denaturation for 2 minutes followed by 33 cycles of 95 °C for 15 s, 55 °C for 15 s and 72 °C for 20 s, ending with one final elongation step at 72 °C for 4 min. The PCR1 products was cleaned using magnetic beads on a Biomek 4000 Workstation © (Beckman Coulter, CA, USA). A second PCR using products from the initial PCR, incorporated standard Nextera Illumina barcodes. 2 μL of initial PCR product plus 4 μL of primer P5 and P7, 5 μL/sample PCRBIO HiFi buffer (PCR Biosystems©, UK) and 0.25 μL/sample PCRBIO HiFI Polymerase (PCR Biosystems ©, UK) was pooled to a total volume of 25 μL. One initial 95 °C denaturation for 1 minute was followed by 13 cycles of 95 °C for 15 s, 55 °C for 15 s and 72 °C for 15 s, ending with one final elongation step at 72 °C for 5 min. The concentration was measured with 1x Qubit dsDNA HS assay Kit (Invitrogen, CA, USA)on Varioskan® (Thermo Fisher Scientific, MA, USA), and finally individual barcodes was added to the samples, before sequencing.

### Microbiota analysis

Quality-control of reads, de-replicating, purging from chimeric reads and constructing zero-radius Operational Taxonomic Units (zOTU) was conducted with the UNOISE pipeline [43] and taxonomically assigned with Sintax [44]. Taxonomical assignments were obtained using the Greengenes (13.8) 16S rRNA gene database. Permutational multivariate ANOVA (PERMANOVA) was used to evaluate group differences based on weighted and unweighted UniFrac distance matrices, or Bray-Curtis dissimilarity metrics, and taxa-level differences were assessed analysis of composition of microbes (ANCOM). Principal Coordinates Analysis (PCoA) was performed on unwehghted UniFrac or Bray-Curtis distances.

### RNA extraction and quantitative PCR

RNA was extracted using a commercial miRNAeasy Mini Kit (Qiagen) following the manufacture’s guidelines. Briefly, tissue was homogenized in Qiazol lysis reagent using a gentleMACS Dissociator (Miltenyi Biotec, Germany) and filtered in an RNAeasy spin column (Qiagen) including on-column DNAase treatment. Afterwards, concentration and purity were measured using a NanoDrop ND-1000 spectrophotometer (NanoDrop Technologies, DE, USA). First-strand cDNA, including gDNA removal, was synthesized using a commercial QuantiTect Reverse Transcription Kit (Qiagen) according to the manufacturer’s instructions. Quantitative PCR was performed using PerfeCTa SYBR Green Fastmix (Quantabio) on a AriaMx Real-time PCR System (Agilent, US under the following conditions: 2 min at 95 °C followed by 40 cycles of 5 sec at 95 °C and 20 sec at 60 °C, and finished with 30 sec at 95 °C, 30 sec at 65 °C and 30 sec of 95 °C again. The primers used were *Dckl1, Duox2, Gpx2, Ifng, Il10 and Gapdh* as reference gene (see Supplementary Table 1 for primer sequences). The CT method was used to calculate fold changes.

### Fluidigm Analysis

A dynamic in-house gut-immunity panel based on genes involved in immunity, gut microbiota signaling and gut barrier functions was used with few modifications from a previous publication [45]. Primers were designed to span an intron if possible and yield products around 75-200 nucleotides long using primer 3 (http://bioinfo.ut.ee/primer3/) or primer blast (https://www.ncbi.nlm.nih.gov/tools/primer-blast/) with standard settings [46, 47]. Primer sequences are listed in Supplementary Table 2. qPCR was performed using the Biomark HD system (Fluidigm Corporation) on a 96.96 IFC using manufacturer’s instructions. The pre-amplification (TaqMan PreAmp, Thermo Fisher Scientific) and the following Exonuclease 1 (NEB Biolabs) treatment were performed on 8x diluted cDNA for 17 cycles using a 250 nM pool of the selected primers. Pre-amplified and exonuclease treated cDNA was diluted 8x before qPCR. Melting curve was assessed in the associated software for multiple peaks and the –RT sample was checked for gDNA background. One primer assay (MVP1) was included as an extra check for gDNA contamination [48]. Primer efficiency was calculated from a calibration curve made from a 5x dilution row of a pool of undiluted pre-amplified and exonuclease treated cDNA. Primer assays with efficiencies between 80-110 % and R >0,98 were accepted for further analysis. All Fluidigm qPCR data processing was performed in Genex6 (multiD Analysis AB). 69 candidate genes and 8 reference genes were assessed. The reference genes were analyzed for stable expression using the geNorm and NormFinder algorithms [49, 50]. *Actb, Sdha, Tbp* and *Ywhaz* were most stable and used as reference genes. Candidate gene expression in cycle of quantification values (Cq) were normalized to the reference genes, cDNA duplicates were averaged, relative expression of the lowest expressed were set to 1 and the data were log2 transformed before statistical analysis.

### ELISA

*T. muris*-specific IgG1, IgG2a and IgE were measured using excretory/secretory antigens and anti-mouse monoclonal antibodies as previously described [30].

### Carbohydrate analyses

Contents of sugars (glucose, fructose, sucrose), fructans, starch, soluble and insoluble non-cellulosic polysaccharides, cellulose, total non-starch polysaccharides, and Klason lignin in the different diets were determined by enzymatic-colorimetric and enzymatic-chemical-gravimetric methods as previously described [51].

### Statistical analysis

Data were analysed using ANOVA or t-tests, or Kruskal-Wallis or Mann-Whitney tests for non-parametric data. Shapiro-Wilk and Kolmogorov-Smirnov tests were used to tests for assumptions of normality in analyses. Data were analyzed using GraphPad Prism 8.3 (GraphPad Software Inc., USA), and significance taken at p < 0.05.

## Acknowledgements

The authors are grateful to Lise-Lotte Christiansen, Mette Schjelde and Denitsa Stefanova for technical assistance, Sophie Stolzenbach for help with DNA extraction, and Dr Sebastian Rausch (Freie Universitat Berlin) and Professor Rick Maizels (University of Glasgow) for provision of *H. polygyrus* larvae and advice and discussions. This work was funded by the Danish Council for Independent Research (Grant 4184-00377).

## Data Availability Statement

Sequence data has been uploaded to NCBI (SRA Bioproject) with the accession number PRJNA821694.

