## Supplementary Table 1 for "Dietary non-starch polysaccharides impair immunity to enteric nematode infection"

**Table S1.** Primers used for qPCR

| **Gene** | **Primer sequence (5'-3' forward/reverse)** |
|---|---|
| *Dclk1* | F: ACTGCAGCAGGAGTTTCTGT<br><br>R: CCGAGTTCAATTCCGGTGGA |
| *Duox2* | F: GGACCACGACAGTGATCTCC<br><br>R: GAACTGTTCCCAGGAGTCCG |
| *Gpx2* | F: TCAAACAGTTCACAGGTGGG<br><br>R: AGTCCTTTAGACCGGTGGGA |
| *Ifng* | F: GACTGTGATTGCGGGGTTGTA<br><br>R: TCACTGCAGCTCTGAATGTTTCT |
| *Il10* | F: GCCCTTTGCTATGGTGTCCT<br><br>R: TAGGGGAACCCTCTGAGCTG |
| *Gapdh* | F: TATGTCGTGGAGTCTACTGGT<br><br>R: GAGTTGTCATATTTCTCGTGG |
