## Supplementary Table 2 for "Dietary non-starch polysaccharides impair immunity to enteric nematode infection"

Primersequences

| Gene name | Forward Primer | Reverse primer | Amplicon | Efficiency% |
|---|---|---|---|---|
| *Actb* | CCCTAAGGCCAACCGTGAAA | CAGCCTGGATGGCTACGTAC | 83 | 109 |
| *Arg1* | ATGGGCAACCTGTGTCCTTT | TCTACGTCTCGCAAGCCAAT | 127 | 98 |
| *Ccl2* | CAGCTCTCTCTTCCTCCACC | TGGGATCATCTTGCTGGTGA | 155 | 110 |
| *Ccl3* | ACCATGACACTCTGCAACCA | CAACGATGAATTGGCGTGGA | 106 | 103 |
| *Ccr10* | AAACCCTTGTAGCCAGAGATGG | CGTACCCGGAGTAAAGTCCC | 70 | 104 |
| *Cd38* | ACTGGAGAGCCTACCACGAA | AGTGGGGCGTAGTCTTCTCT | 179 | 104 |
| *Cd3e* | CCTCCTAGCTGTTGGCACTT | GGGCACGTCAACTCTACACT | 90 | 107 |
| *Cd8a* | GGATTGGACTTCGCCTGTGA | TGGGACATTTGCAAACACGC | 130 | 94 |
| *Cdh1* | GAGACCAGTTTCCTCGTCCG | AGCAGCTCTGGGTTGGATTC | 137 | 105 |
| *Cgn* | TGAATCCGGGAGCACTGATCT | ACGAAGGGTGCCCATTTCAG | 151 | 106 |
| *Cldn1* | TCGACTCCTTGCTGAATCTGA | CAGCCATCCACATCTTCTGC | 159 | 109 |
| *Cldn7* | GCCTTGGTAGCATGTTCCTG | TTTGCTTTCACTGCCTGGAC | 180 | 110 |
| *Csf2* | ATGCCTGTCACGTTGAATGA | CCGTAGACCCTGCTCGAATA | 108 | 85 |
| *Ctla4* | ATGGCTTGTCTTGGACTCCG | ACCACTGAAGGTTGGGTCAC | 137 | 103 |
| *Ctnnb1* | GAGCACATCAGGACACCCAA | CCGAGCAAGGATGTGGAGAG | 122 | 108 |
| *Cxcl1* | TGCACCCAAACCGAAGTCAT | CTCCGTTACTTGGGGACACC | 122 | 105 |
| *Cxcl16* | CCCAGATACCGCAGGGTACTT | TTCCCATGACCAGTTCCACA | 181 | 109 |
| *Cxcl10* | AAGTGCTGCCGTCATTTTCT | CCTATGGCCCTCATTCTCAC | 129 | 101 |
| *Cxcr6* | TGGAACAAAGCTACTGGGCT | TCGTAGTGCCCATCGTACAG | 81 | 92 |
| *Defa3* | AAAACTGAGGAGCAGCCAGG | CAGCGACAGCAGAGTGTGTA | 194 | 96 |
| *Fasl* | AAGGAACTGGCAGAACTCCG | ACTCCAGAGATCAGAGCGGT | 151 | 90 |
| *Ffar1* | CTGGGCATCAACATACCCGT | AGCAGAAGGCAGTGATGACC | 133 | 92 |
| *Ffar4* | TGCCCCTCTGCATCTTGTTC | GGTTGGGCCAATCCAATGTG | 90 | 106 |
| *Foxp3* | AGAGAGAAGTGGTGCAGTCTC | GAGTACTGGTGGCTACGATG | 159 | 97 |
| *Fxr* | GCTGAGACTGGGTACCAGGG | TCGGAAGAAACCTTTGCAGCC | 183 | 105 |
| *Gata3* | GCTACGGTGCAGAGGTATCC | CAGAGATCCGTGCAGCAGAG | 75 | 110 |
| *Gcg* | TCTACACCTGTTCGCAGCTC | GTCCTCATGCGCTTCTGTCT | 172 | 103 |
| *Hcar2* | CTTCCTACCCAGTGTGGCTG | CAGGTCCACCGAGGAGTAGA | 94 | 106 |
| *Hprt* | TCAGTCAACGGGGGACATAAA | GGGGCTGTACTGCTTAACCAG | 122 | 86 |
| *Icam1* | CTGTGCTTTGAGAACTGTGGC | CAGGGTGAGGTCCTTGCCTA | 129 | 101 |
| *Ifng* | TTTGAGGTCAACAACCCACAG | GCTTCCTGAGGCTGGATTC | 94 | 96 |
| *Il10* | AGGCGCTGTCATCGATTTCT | ATGGCCTTGTAGACACCTTGG | 104 | 108 |
| *Il12b* | TTGTTCGAATCCAGCGCAAG | TTCTCTACGAGGAACGCACC | 83 | 105 |
| *Il15* | ACAGCTCAGAGAGAATCCACC | ATGAGCTGGCTATGGCGATG | 187 | 89 |
| *Il17a* | TGAGTCCAGGGAGAGCTTCA | CGCTGCTGCCTTCACTGTA | 80 | 89 |
| *Il1b* | GCAACTGTTCCTGAACTCAACT | ATCTTTTGGGGTCCGTCAACT | 89 | 100 |
| *Il2* | GAAACTCCCCAGGATGCTCA | CGCAGAGGTCCAAGTTCATCT | 99 | 88 |
| *Il27* | GTCCACAGCTTTGCTGAATCT | CGAAGTGTGGTAGCGAGGAA | 149 | 99 |
| *Il33* | GGGCTCACTGCAGGAAAGTA | TTTGCCGGGGAAATCTTGGA | 115 | 101 |
| *Il4* | CCTGGATTCATCGATAAGCTG | TCCATTTGCATGATGCTCTT | 93 | 99 |
| *Il6* | GACAAAGCCAGAGTCCTTCAGA | AGGAGAGCATTGGAAATTGGGG | 113 | 102 |
| *Irf3* | CCACAAGGACAAGGACGGAG | CCACATTTCCCCCATGCAGA | 124 | 101 |
| *Itgax* | GAGCCAGAACTTCCCAACTG | ACCCGAGCCATCAATCAGG | 79 | 96 |
| *Muc1* | AGTACCAAGCGTAGCCCCTA | GTGGGGTGACTTGCTCCTAC | 118 | 108 |
| *Muc2* | TATGCCAGGCCAGGAGTTTA | GCAAGGCAGGTCTTTACACA | 82 | 101 |
| *Muc4* | GTCCACTTCTTCCCCATCTCG | CCATTGTGACAGTAGCCCTCA | 173 | 103 |
| *MVP1* | GGAGCCCAGTGTAGAAGAGCA | AGCCAGCGAACCATATCCTGA | 87 | - |
| *Myd88* | CCAGGTGTCCAACAGAAGC | CTTGGTGCAAGGGTTGGTAT | 114 | 101 |
| *Nfkb* | GGCAGGTATTTGACATACTAAATGG | TGCAGAGTTGTAGCCTCGTG | 117 | 102 |
| *Nfkbia* | GAGCGAGGATGAGGAGAGCTA | GGCCTCCAAACACACAGTCA | 83 | 107 |
| *Nod2* | TGGCCCTACAGCTGGATTAC | TTGTTGTTGAAGAGACTGGCTA | 187 | 110 |
| *Ocld* | GCTGCTGCTGATGAATATAATAGACT | TCCCACCATCCTCTTGATGT | 120 | 104 |
| *Pgk1* | GGTGTTGCCAAAATGTCGCT | GGACTTGGCTCCATTGTCCA | 183 | 109 |
| *Pla2g2a* | GGGGCCAAATCACCTGTTCT | GTTCCGGGCGAAACATTCAG | 92 | 105 |
| *Ppara* | AACATCGAGTGTCGAATATGTGG | CCGAATAGTTCGCCGAAAGAA | 99 | 103 |
| *Pparg* | TTCAGAAGTGCCTTGCTGTG | CCAACAGCTTCTCCTTCTCG | 84 | 105 |
| *Ppia* | CCACCGTGTTCTTCGACATC | AGTGCTCAGAGCTCGAAAGT | 113 | 112 |
| *Prkaa2* | GCAAAGTGAAGACTACCAGGTG | GTAATCCACGGCAGACAGGA | 163 | 110 |
| *Pyy* | GCTTCTCCCACCTTCCATCT | AGACAGGCGAGCAGGATTAG | 121 | 106 |
| *Reg3g_v2* | CCCTCAGGACATCTTGTGTCT | ACCTCTGTTGGGTTCATAGCC | 141 | 103 |
| *Reg3g* | ACAGACAAGATGCTTCCCCG | AGCTGCTACGTGAAGATGGG | 127 | 105 |
| *Retnlb* | CTGTCCTGCTGGGATGGT | CCAGTCCATGACTGAGCACT | 109 | 98 |
| *Sdha* | ATTGCTACTGGGGGCTACGG | GTCCTGGCAAGGCAAACCAG | 108 | 106 |
| *Stat4* | GAAGTACCTCTACCCTGACATTCC | AGGGGACGTAACCCTTGTCT | 113 | 99 |
| *Stat5* | GGTCCCTGAGTTCGTCAATG | GGTTGGGTGGGTACATGTTG | 116 | 110 |
| *Tbp* | ACCTTATGCTCAGGGCTTGG | TGCCGTAAGGCATCATTGGA | 83 | 88 |
| *Tbx21* | GGGCTTCCAACAATGTGACC | AGCTGAGTGATCTCTGCGTTC | 193 | 105 |
| *Tff3* | CTGTCACATCGGAGCAGTGT | CAGGGCACATTTGGGATACT | 67 | 107 |
| *Timp1* | GGGGTGTGCACAGTGTTTC | GACCTGATCCGTCCACAAAC | 81 | 107 |
| *Tlr2* | GCATCCGAATTGCATCACCG | ACAGCGTTTGCTGAAGAGGA | 136 | 99 |
| *Tlr3* | GAATCACAATCGCGCACCAA | CCATAGGGACAAAAGTCCCCC | 178 | 85 |
| *Tlr4* | CTCTCATGGCCTCCACTGGT | TTAGGAACTACCTCTATGCAGGGAT | 137 | 104 |
| *Tlr5* | GATGGATGCTGAGTTCCCCC | AAAGGCTATCCTGCCGTCTG | 139 | 91 |
| *Tnfa* | CAAATGGCCTCCCTCTCATCA | TGGGCTACAGGCTTGTCAC | 88 | 110 |
| *Tnfsf15* | CCATCCTCGCAGGACTTAGC | TGCCTCTGGGAGGTGAGTAA | 135 | 101 |
| *Tuba* | TGTCCTGGACAGGATTCGC | CTCCATCAGCAGGGAGGTG | 115 | 108 |
| *Ywhaz* | GAAAAGTTCTTGATCCCCAATGC | TGTGACTGGTCCACAATTCCTT | 134 | 111 |
| *Zbtb16* | GCACTACAGGGTTCACACAGG | CACCGTTGTGTGTTCTCAGG | 107 | 104 |
